# MRI-free Virtual Neuronavigation for TMS

**DOI:** 10.1101/2025.04.10.648190

**Authors:** Yishai Valter, Dennis Q Truong, Yushi Kawasumi, Marom Bikson, Abhishek Datta

## Abstract

Methods to determine coil position are paramount in TMS. Existing approaches are limited to scalp heuristics or depend on some combination of neuronavigation hardware, average brain templates, as well as subject MRI and/or evoked responses. We developed head-model guided TMS virtual neuronavigation without subject MRI, specialized hardware, or evoked responses. The MRI-free virtual neuronavigation involves three scalp measurements which are then used to generate an individualized head model by applying ellipsoid-based affine transformations to the MNI standard head model. Virtual neuronavigation is then performed on this individualized head model for any brain target. The coil position is then provided to the operator in simple geodesic measurements. We simulated this process on MRI data of fifteen subjects showing a mean error of 2.75 mm, outperforming the accuracy of scalp heuristic targeting approaches.

## Introduction

All repetitive transcranial magnetic stimulation (rTMS), for investigational or clinically-approved treatment (Cohen et al., 2022), requires a method to position a coil on the scalp. rTMS coils are positioned according to application-specific cortical targets, such as the left dorsolateral prefrontal cortex (DLPFC) for depression treatment. Since anatomy varies, coil placement relies on varied approaches that are head-model based (1-3) or head-model absent (4-5):

Head-model based:

1. MRI-Guided Neuronavigation: The gold-standard approach uses individual magnetic resonance imaging (MRI) and stereotactic neuronavigation hardware to co-register the coil with the subject’s MRI in real time (Neggers et al., 2002; Comeau, 2014; Ruohonen et al., 2010; Valter et al., 2024).
2. Template-Based Neuronavigation: When individualized MRI data is unavailable, stereotactic navigation hardware can be used to map anatomical landmarks in 3D space. Algorithms then warp a standard head model, such as the Montreal Neurological Institute template (MNI152; Mazziotta et al., 2001) to subject-specific head size and shape (Valiulis et al., 2019). Brain targets are approximated via affine transformations, and the coil is guided by neuronavigation hardware accordingly (Fleischmann et al., 2020; Ross et al., 2022; Valiuliene et al., 2023).
3. Virtual Neuronavigation: When an individual MRI is available but neuronavigation hardware is not, virtual neuronavigation identifies the target location and prescribes coil placement using individualized geodesic measurements or customized headgear (Mir-Moghtadaei et al., 2015; Vaghefi et al., 2015; Wang et al., Hu 2024). Head-model absent:
4. Generic geodesic (Scalp Heuristics): Due to the cost and complexity of navigation systems, localization based on scalp measurements using external landmarks are common. The Beam-F3 method is widely used for DLPFC targeting (Beam et al., 2009; McClintock et al. 2017). However, concerns about its accuracy have led to the development of alternative geodesic measurement techniques based on group averages (Fabregat-Sanjuan et al., 2022; Mir- Moghtadaei et al., 2022; Jiang et al., 2022; Li et al., 2025).
5. Relative to TMS evoked responses: Approaches like the “5-cm rule” combine a functional evoked response (e.g. TMS-MEP) with a geodesic rule (e.g. “5 cm”) (O’Reardon et al., 2007).

We developed an approach for TMS coil placement that is head-model based, allowing navigation to any target, but does not require an individual MRI or neuronavigation hardware. This approach combines minimal cost, simple set-up, with virtual head-model based targeting. Our method relies solely on scalp tape-measurements, used in a novel approximation of head dimensions. Then, an individualized head model is generated by applying an affine transformation to the MNI152 template in accordance with the subject’s head geometry and size. Any scalp target can then be identified based on MNI coordinates. Finally, the individuated coil position is prescribed using simple geodesic measurements.

Validation of our head model was performed on a cohort of fifteen healthy volunteers by comparing the pipeline-predicted scalp dimensions with the subjects’ empirically measured scalp dimensions. We subsequently demonstrated that the coil positioning for a DLPFC target achieves an error of less than 3 mm relative to the gold-standard placement, outperforming conventional scalp-based heuristic methods.

## Methods

There is an intrinsic challenge in obtaining the linear dimensions of a round, solid object when physical tools like a tape measure cannot traverse its interior (e.g., through the head). While large calipers can yield dimensions for scalp length and width, they cannot be used to measure scalp height and precise versions are not practical in clinical practice. Conversely, a tape measure is far more accessible and is routinely used in rTMS administration.

While concentric sphere head models have been used extensively for brain stimulation modelling (Datta et al., 2008; Deng et al., 2013) and EEG electrode positioning systems (Cobb et al., 1958, Oostenveld & Praamstra, 2001), visual observation suggests that an ellipsoid offers improved anatomical fidelity by accommodating variations in scalp dimensions that a uniform spherical model cannot capture. Indeed, ellipsoid conceptualizations of the head have been used to model mechanical impact (Heydari & Jani, 2010) and auditory processing (Duda et al., 1999). Given the framework for an ellipsoid-based head model but without a practical method to measure ellipsoid linear dimensions—we developed a mathematical formula to calculate the semi-primary axes (radii) of an ellipsoid from circumferential measurements. We hypothesized that this formula would enable accurate approximation of scalp dimensions using only a tape measure.

### Ellipsoid Formula

The general equation of an ellipsoid is 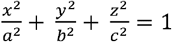 where *a,b*, and *c* are the ellipsoid’s three axes. When *x,y*, or *z* are set to 0, the ellipsoid equation simplifies to either one of the following 2- dimensional ellipses

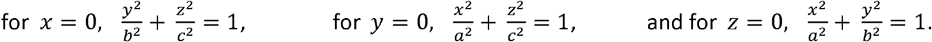

Importantly, the axes of each of these ellipses are the same as the parent ellipsoid. Calculating the axes of the parent ellipsoid is therefore possible by calculating the axes of each independent ellipse, which can be done using Ramanujan’s commonly used approximation equation for the perimeter of an ellipse (Ramanujan, 1914; Villarino, 2005). For an ellipse with perimeter *p* and axes *a* and *b*, Ramanujan’s equation states

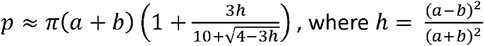

Applying this equation, a system of three equations can be created, one equation for each ellipse:

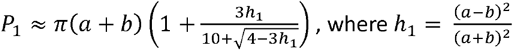

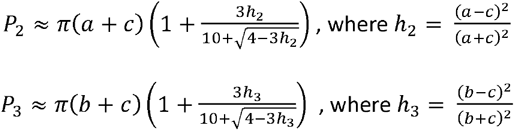

When *P*_1_, *P*, and *P*_3_ are empirically obtained, this system of non-linear equations can be solved using numerical methods, such as with the `vpasolve()` function in MATLAB, to obtain the values of *a, b*, and *c*.

### MRI-free Head Modeling

According to the 10-10 EEG system, the scalp can be circumscribed by an ellipse that intersects the Fpz and Oz positions (Jurcak et al., 2007). Using this map, three circumferential measurements (*P*_l_, *P*_2_, and *P*_3_) can be obtained using a tape measure (**Figure 1A**). For *P*_l_, we propose measuring the full head circumference at Fpz-Oz height. Then, locate the midpoints between Fpz and Oz on either side of the head to serve as the lateral extrema of the scalp (approximately T3 and T4 in the EEG system). Measure the geodesic distance between these two midpoints passing over the vertex (Cz). This distace is then doubled to obtain *P*_2_, to estimate the entire ellipse perimeter, rather than just the top hemisphere. For *P*_3_, measure the geodesic distance from Fpz to Oz passing over Cz. The distance is also then doubled to estimate the entire ellipse perimeter, rather than just the top hemisphere. Importantly, all three of these measurements can be obtained using only a simple tape measure without the need for any advanced equipment or training. The measured values for *P*_l_, *P*_2_, and *P*_3_ can then be used to approximate the scalp’s radii by applying the equations presented earlier for calculating a, b, and c. Once the dimensions of a subject’s head have been approximated, a scaling matrix can be applied to the MNI152 template to transform it to the subject’s individual dimensions. The head model generated can then be segmented into different anatomical structures, converted into a 3D mesh, and used for a variety of applications, such as virtual neuronavigation.

**Figure 1.**
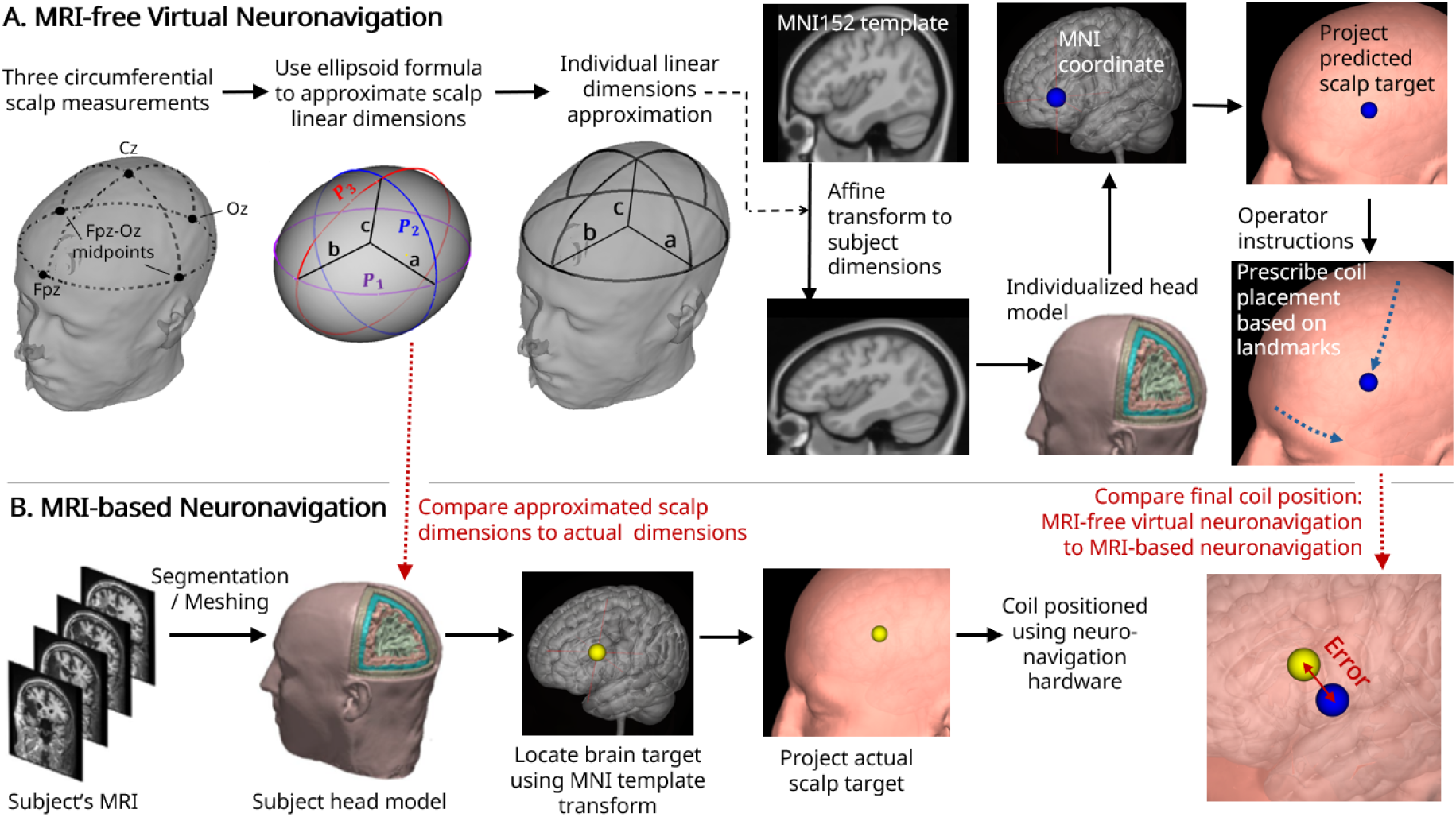
A. MRI-free Virtual Neuronavigation: Three circumferential measurements are performed and used to approximate the scalp’s linear dimensions using an ellipsoidal model. An affine transformation is then applied to the MNI152 template to warp it to the subject’s scalp dimensions. Subsequently, a brain target is identified in the individualized head model and projected to the scalp. Virtual neuronavigation is then performed on the individualized head model to provide operator instructions to locate the scalp target. **2. MRI-based Neuronavigation:** For this targeting method, subject-specific MRI is used to identify the brain and scalp targets. A TMS coil is then positioned at the target using neuronavigation hardware. The Euclidean distance between the two targets is the error of MRI-free Virtual

### Data Acquisition and Pre-processing

We obtained structural MRI of 15 adult subjects (4 female) from an ongoing trial. MRI were acquired on a Siemens 3T Prisma scanner equipped with a 32-channel head coil and an imaging sequence including T1-weighted MPR vNav scans at 0.8 mm isotropic. The subject races were Asian (5) and White or Caucasian (10), providing a variety of head shapes. The study was approved by the WCG IRB (Study No. 1343183) and performed in accordance with the Declaration of Helsinki. Subjects provided written consent before scanning. The study was registered on clinicaltrial.gov (No. NCT05598931). After MRI acquisition we automated segmentation and meshing using the CHARM command included in the SimNIBS open-source software package v4.1.0 (Thielscher et al., 2015; Saturnino et al., 2019; Puonti et al., 2020). The CHARM pipeline also generates a csv file with real-world coordinates for electrode locations on the subject’s scalp according to the 10-10 EEG system of Jurcak et al. (2007). We used these coordinates for all scalp measurements.

### Scalp Measurements

We performed scalp measurements *in silico* using the curvilinear measurement tool in the commercial software OsiriX v14 (Prixmeo SARLA, Bernex, Switzerland). *In silico* scalp measurements are generally considered analogous to *in vivo* measurements with a cap, pen, and tape measure in clinical practice (Li et al., 2025). We then approximated the scalp radii of the 15 subjects from these scalp measurements by applying the ellipsoid formula presented earlier. We also empirically obtained the actual scalp radii, where radius ‘a’ is the Euclidean distance between the two lateral scalp extrema (left-to-right midpoints) divided by two, radius ‘b’ is the Euclidean distance between Fpz and Oz divided by two, and radius ‘c’ is the Euclidean distance between the Fpz-Oz centroid and Cz. We compared the predicted radii to the true radii of each subject’s scalp to test the accuracy of our approach (**Figure 1B**).

### Transforming MNI152 Template to Subject Dimensions

Transformations were applied to the MNI152 template (Non-linear 6th Generation) using 3D Slicer v5.6.2, an open-source platform for medical imaging analysis (Fedorov et al., 2012). To ensure accurate scaling, the template was first centered using a translation vector of [-0.625, 15.35, -4.25] and rotated 5° clockwise in the sagittal plane (y-z plane) to align the Fpz-Oz line parallel to the y-axis. Following this adjustment, the template was scaled to match each subject’s estimated dimensions using the scaling matrix:

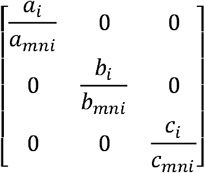

Where *a*_*i*_,*b*_*i*_,*c*_*i*_ are each subject’s scalp dimensions and *a*_*mni*_, *b*_*mni*_, *c*_*mni*_ are the MNI152 head dimensions which we had previously computed to be 10.29 cm, 8.38 cm, and 9.33 cm respectively.

### Identifying Brain and Scalp Target

For demonstration, we selected the MNI coordinates proposed by Fox et al. (2014) for TMS depression treatment [-38, 44, 26] as previously selected to test the concordance of Beam-F3 with MRI-guided neuronavigation (Mir-Moghtadaei et al., 2015). Normalization to MNI space was done using the maff8 function in Brainstorm, a Statistical Parametric Mapping mutual information algorithm that implements a 4×4 linear transformation (Friston et al., 1995; Ashburner & Friston, 2005). To identify the corresponding scalp location for the MNI brain coordinate, we used the ‘Project to Scalp’ tool in SimNIBS which applies an expanding sphere at the brain coordinate to the point of intersection of the expanding sphere with the scalp surface. The point of intersection is the nearest scalp point to the brain target.

### MRI-free Virtual Neuronavigation

The individualized MNI images were imported into OsiriX where the geodesic distances between the scalp target and landmarks were measured following the method of Mir-Moghtadaei et al. (2015). Specifically, we first measured the geodesic distance between the vertex and scalp target and then continued the line until it intersected the Fpz-Oz circumferential line (“Intersection Point”). We then measured the geodesic distance from Intersection Point to Fpz. This produced two scalp vectors, akin to the Beam-F3 approach, which could then be used by a clinician to locate the scalp target without requiring a neuronavigation system. We then simulated the actions of a clinician by replicating these geodesic instructions on the subjects’ MRI models.

### Accuracy Testing of Virtual Neuronavigation

In parallel, we also identified the actual ideal scalp target using the subjects’ MRI data via spatial normalization and brain-to-scalp projection as was done for the individualized MNI heads using the maff8 transformation and SimNIBS. This provided the actual scalp location on the subjects’ heads corresponding to the ideal MNI brain coordinates for depression treatment. Determining the Euclidean distance between the empirically obtained scalp target and the spot reached by measuring the recorded distances on the subjects’ heads served to test the accuracy of our head modeling and virtual neuronavigation workflow.

## Results

### Scalp Dimensions

As shown in **Figure 2**, we found a high congruence between estimated and observed scalp dimensions with mean error magnitudes of 1.68% (SD ± 1.12%) in scalp width, 1.65% (SD ± 1.02) in scalp length, and 1.66% in scalp height (SD ± 0.98%). Noticeably, scalp length was consistently over-approximated, possibly due to the spherical shape of the posterior scalp that extends its circumference beyond that of an ellipsoid. This discrepancy may cause the measured perimeters to exceed the corresponding dimension of a perfect ellipsoid. Post-hoc analysis indicated that this systematic offset can be mitigated by applying a 1% adjustment factor to the approximated length (i.e., multiplying the predicted value by 0.99). Incorporating this correction reduces the mean absolute error magnitude in length from 1.65% to 0.88%.

**Figure 2:**
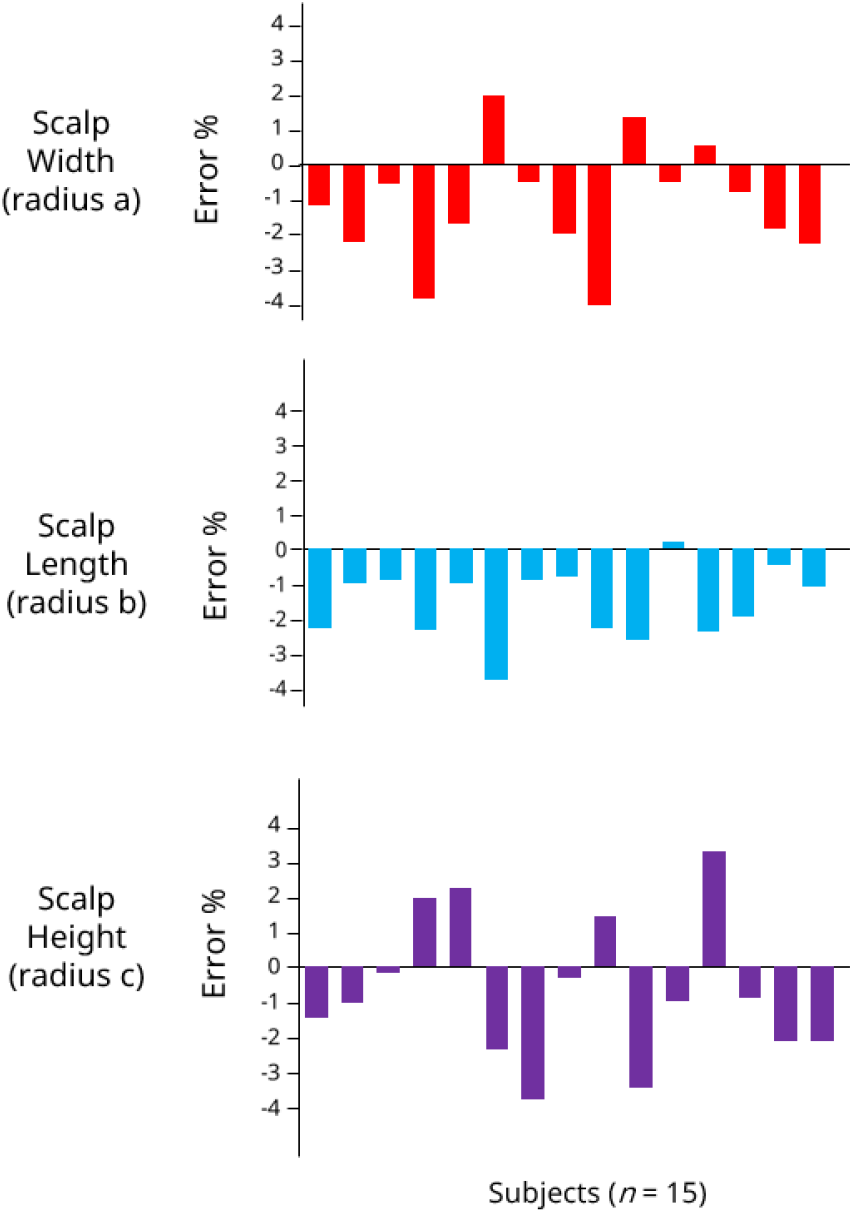
Error percentages between predicted scalp dimensions and actual scalp dimensions.

### Head modeling and Neuronavigation

After applying the 1% adjustment factor, we transformed the MNI152 template to each of the 15 subjects’ scalp dimensions. We then performed virtual neuronavigation on each generated head model to obtain geodesic measurements for locating the specific MNI coordinates associated with depression treatment as done by Mir-Moghtadaei et al. (2015). As shown in **Figure 3**, the mean error between the virtually navigated scalp target and the actual scalp target was 2.75 mm ± 1.46 mm, demonstrating greater accuracy than scalp-heuristic approaches.

**Figure 3:**
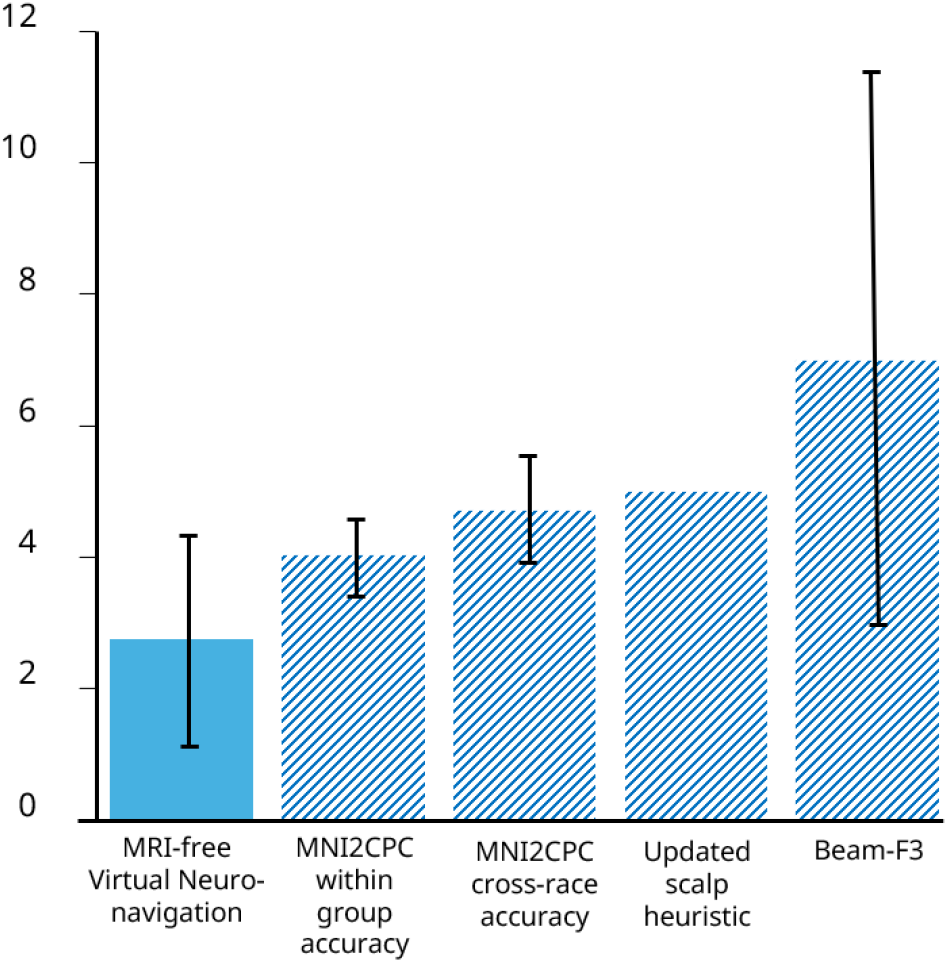
Mean errors (mm) ± SD for MRI-free virtual neuronavigation compared to previously reported scalp heuristics. Jiang et al., (2022) developed the MNI2CPC method and report within-group accuracy (Chinese subjects) as well as the accuracy when applying their method on Caucasian subjects (cross-race accuracy). They also tested the Updated Scalp Heuristic developed by Mir-Moghtadaei et al. (2022) on four scalp targets and reported the error for each target. We provide the mean of the four reported errors. The discrepancy between Beam- F3 and optimal MNI coordinates for depression treatment was reported

## Discussion

Compared to other TMS coil positioning techniques, MRI-free Virtual Neuronavigation for TMS offers specific advantages. It does not require MRI or neuronavigation hardware, yet yields an individualized head model that can be used to guide a coil to any MNI-based brain target with high accuracy. Our approach can therefore be integrated into existing TMS practices without increasing cost or time.

Our platform lends itself to further refinements including 1) use of race-specific head templates (Bhalerao et al., 2018; Yang et al., 2020) to accommodate group-mediated differences; 2) more anatomically faithful representation of the head anatomy could be achieved by decomposing its structure into a combination of multiple geometric elements rather than a single ellipsoid; 3) refined transformation algorithms using different anatomical reference points either in combination with or instead of 10-10 EEG points, thereby enhancing geometric fidelity; 4) integration with low-cost imaging technologies to improve accuracy while maintaining accessibility; 5) integration of electric field models; 6) AI-based method for generating high-resolution MR images (Juan et al., 2023).

While our approach was initially demonstrated for TMS, it is inherently generic and holds promise for a range of neuromodulation modalities. By facilitating the generation of a comprehensive head model derived solely from scalp measurements, our method exhibits potential for targeting deep brain structures using modalities such as transcranial temporal interference stimulation (Grossman et al., 2017; Huang et al., 2020) and transcranial focused ultrasound (Lee et al., 2022; Truong et al., 2022). Future research is necessary to validate the efficacy and reliability of our approach across these applications. Moreover, the versatility of our modeling technique may extend beyond neuromodulation to encompass diverse applications ranging from surgical planning (Singh & Singh, 2021) and anthropometric analysis (Farkas et al., 1992) to the design and evaluation of customized headgear (Liu et al., 2008; Lacko et al., 2017).

## Declarations

### Availability of Data and Materials

The raw data used for this study is available to the public upon reasonable request from the authors.

### Author’s Contributions

Y.V. conceptualization, methodology, data analysis, initial draft; D.Q.T. data curation; M.B. conceptualization, methodology; A.D. funding acquisition, methodology; All authors reviewed and edited the manuscript.

### Generative AI Statement

During the preparation of this work the authors used generative AI in order to improve language and readability. After using generative AI, the authors reviewed and edited the content as needed and take full responsibility for the content of the publication.

